# Spatial vision by macaque midbrain

**DOI:** 10.1101/171645

**Authors:** Chih-Yang Chen, Lukas Sonnenberg, Simone Weller, Thede Witschel, Ziad M. Hafed

## Abstract

Visual brain areas exhibit tuning characteristics that are well suited for image statistics present in our natural environment. However, visual sensation is an active process, and if there are any brain areas that ought to be particularly “in tune” with natural scene statistics, it would be sensory-motor areas critical for guiding behavior. Here we found that the primate superior colliculus, a structure instrumental for rapid visual exploration with saccades, detects low spatial frequencies, which are the most prevalent in natural scenes, much more rapidly than high spatial frequencies. Importantly, this accelerated detection happens independently of whether a neuron is more or less sensitive to low spatial frequencies to begin with. At the population level, the superior colliculus additionally over-represents low spatial frequencies in neural response sensitivity, even at near-foveal eccentricities. Thus, the superior colliculus possesses both temporal and response gain mechanisms for efficient gaze realignment in low-spatial-frequency dominated natural environments.

## Introduction

The superior colliculus (SC) is a visual-motor structure important for transforming visual signals into behaviorally-appropriate gaze shift commands (Boehnke and Munoz, 2008; Gandhi and Katnani, 2011; Optican, 2005; Sparks and Mays, 1990; Veale et al., 2017; Wurtz, 1996). Even though much is known about the SC's afferent and efferent connections, as well as its physiological visual and eye-movement-related neural response characteristics, such knowledge has predominantly been obtained using highly impoverished stimuli, like small spots of light presented over an otherwise uniform background. However, ecological constraints (Hafed and Chen, 2016; Previc, 1990) on both visual perception and eye movements imply that the SC, like other brain regions, should best function if its neurons’ properties were well matched with the properties of the environment.

Among such properties is the preponderance of low spatial frequencies in natural scene statistics (Ruderman and Bialek, 1994; Tolhurst et al., 1992). In early visual areas, such preponderance is well matched with a variety of observations, including coarse-to-fine neural image analysis (Bredfeldt and Ringach, 2002; Mazer et al., 2002; Purushothaman et al., 2014) and neural image filtering kernels that are suitable for natural scene statistics (Olshausen and Field, 1996; Simoncelli and Olshausen, 2001; van Hateren and van der Schaaf, 1998). Curiously, such observations are often also used to account for motor rather than perceptual effects, for example on manual and saccadic reaction times (Breitmeyer, 1975; Ludwig et al., 2004; White et al., 2008), even though these early visual areas may be viewed as being more relevant for perception rather than action.

In this study, we hypothesized that the SC's importance in guiding action (Gandhi and Katnani, 2011; Veale et al., 2017) should make it as well matched to spatial properties present in natural scenes as early visual areas, if not more so, and in a manner that is highly conducive of behavioral motor effects. We specifically tested the ability of SC neurons to detect low spatial frequency visual stimuli. We found that these neurons do so much earlier than for high spatial frequencies, and independently of neural sensitivity to a given spatial frequency. Moreover, we found that at the population level, SC neural sensitivity to spatial frequency was primarily low-pass in nature, meaning that both SC response time and SC response strength are particularly efficient when visually analyzing the low spatial frequencies that are abundantly present in natural scenes. These observations have allowed us to predict, with high fidelity, our animals’ saccadic reaction time patterns as a function of spatial frequency based solely on SC visual response strength and latency measurements obtained from completely different experimental sessions not involving saccadic responses. We believe that our findings clarify important visual functions of the SC, complementary to this structure's more well-studied motor (Gandhi and Katnani, 2011) and cognitive (Krauzlis et al., 2013) functions. Such findings not only allow better understanding of visual-motor behavior under more naturalistic conditions than with impoverished laboratory stimuli, but they also help clarify potential underlying mechanisms for pathological cases in which the SC's role in vision may be magnified. For example, in blindsight (Weiskrantz et al., 1974), patients with V1 loss exhibit spatial frequency tuning properties remarkably similar to those that we describe here (Sahraie et al., 2010; Sahraie et al., 2002; Trevethan and Sahraie, 2003).

## Results

### Faster SC responses to low spatial frequencies irrespective of neural sensitivity

We recorded visual responses in macaque monkeys that were passively fixating a small spot of light (Chen and Hafed, 2013; Chen et al., 2015). During such passive fixation, we presented a high contrast sine wave grating filling the visual response field (RF) of a recorded neuron (Materials and Methods). We randomly varied the spatial frequency of the presented grating from trial to trial and noticed a systematic rank ordering of neural response latencies as a function of spatial frequency. For example, in the neuron depicted in Fig. 1A, visually-evoked action potentials arrived earliest for gratings of 0.56 or 1.11 cycles/deg (cpd), and their latency progressively increased for higher spatial frequencies. This observation is reminiscent of coarse-to-fine image coding properties of early cortical visual areas (Bredfeldt and Ringach, 2002; Mazer et al., 2002; Purushothaman et al., 2014), but it still violated an expected inverse relationship between response latency and response sensitivity (i.e. response gain) that has been reported in both the SC (Marino et al., 2012) and in early cortical visual areas (Maunsell and Gibson, 1992). Specifically, visual sensitivity in this neuron was highest for 4.44 cpd (Fig. 1B), but visual response latency at this spatial frequency was significantly longer than at lower frequencies (first-spike latency at 4.44 cpd was 74.61 ms +/- 0.42 ms s.e.m.; first-spike latency at 0.56 cpd was 51.49 ms +/− 0.82 ms s.e.m.; p=1.14×10^−38^, Ranksum test). This meant that plotting tuning curves of either visual sensitivity (Fig. 1C, top; Materials and Methods) or visual latency (Fig. 1C, bottom) as a function of spatial frequency revealed a dissociation between the two neural response properties: the preferred spatial frequency in terms of response sensitivity was ~4 cpd, whereas the preferred spatial frequency in terms of response latency was much lower.

**Figure 1.**
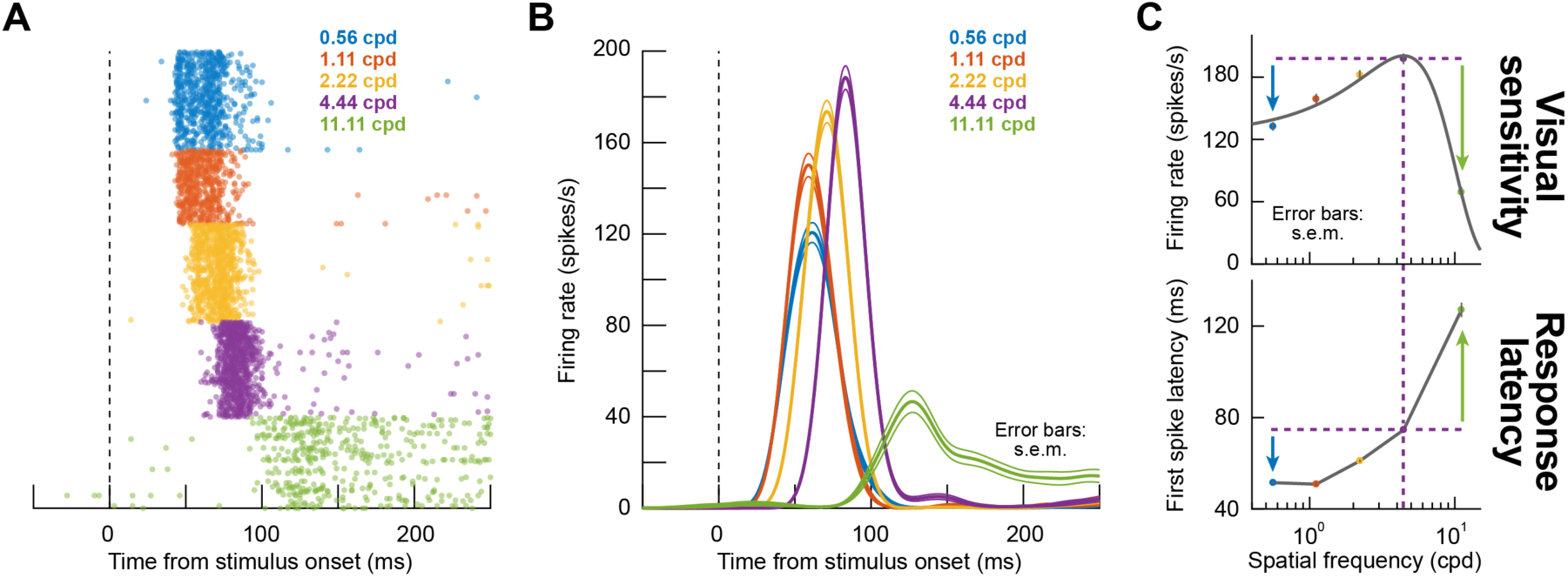
Rapid detection of low spatial frequencies by macaque superior colliculus (SC). (**A**) Visual responses of an example SC neuron to gratings of different spatial frequencies (color-coded according to the legend). The raster plots show times of individual action potentials with different trials stacked in rows, and the trials were grouped by color only for presentation in the figure; in the experiment, different spatial frequencies were presented randomly. There was a rank ordering of response latency, with the lowest spatial frequencies (e.g. 0.56 and 1.11 cpd) evoking the shortest-latency neural responses. (**B**) This effect happened even though higher spatial frequencies (e.g. 2.22 and 4.44 cpd) elicited stronger responses. This is illustrated by plotting firing rates for the same neuron, since it is easier to infer response amplitudes from firing rates. The neuron emitted the strongest visual responses for 4.44 cpd gratings even though these strong responses came later than for lower spatial frequencies. Thus, there was a dissociation between response latency and response sensitivity. (**C**) Tuning curves illustrating the dissociation. The top panel plots the tuning curve of the neuron according to response sensitivity (i.e. response amplitude as a function of spatial frequency; Materials and Methods). Visual responses were strongest for 4.44 cpd gratings with both lower and higher spatial frequencies (e.g. the colored arrows) evoking significantly weaker responses. On the other hand, in the lower panel, the latency to first visually-evoked spike (Materials and Methods) at 4.44 cpd was longer than for lower spatial frequencies but shorter than for higher spatial frequencies (e.g. the colored arrows). Error bars in **B**, **C**, when visible, denote s.e.m. across trials.

We confirmed the dissociation between visual sensitivity and visual latency across our recorded population. For example, for neurons preferring 4.44 cpd in terms of visual sensitivity, we plotted either such sensitivity (Fig. 2A) or instead response latency (Fig. 2B) for different spatial frequencies; in all cases, we compared responses to those obtained when the preferred 4.44 cpd gratings were presented. For example, in the leftmost panel of Fig. 2A, we plotted response sensitivity to 0.56 cpd (bluish dots) or 11.11 cpd (greenish dots) as a function of response sensitivity to 4.44 cpd. Since the neurons preferred 4.44 cpd, response sensitivity was naturally lower for both 0.56 cpd and 11.11 cpd (p-values for statistical tests are shown in the figure). Similar results were obtained in the middle and rightmost panels of Fig. 2A for other spatial frequencies.

**Figure 2.**
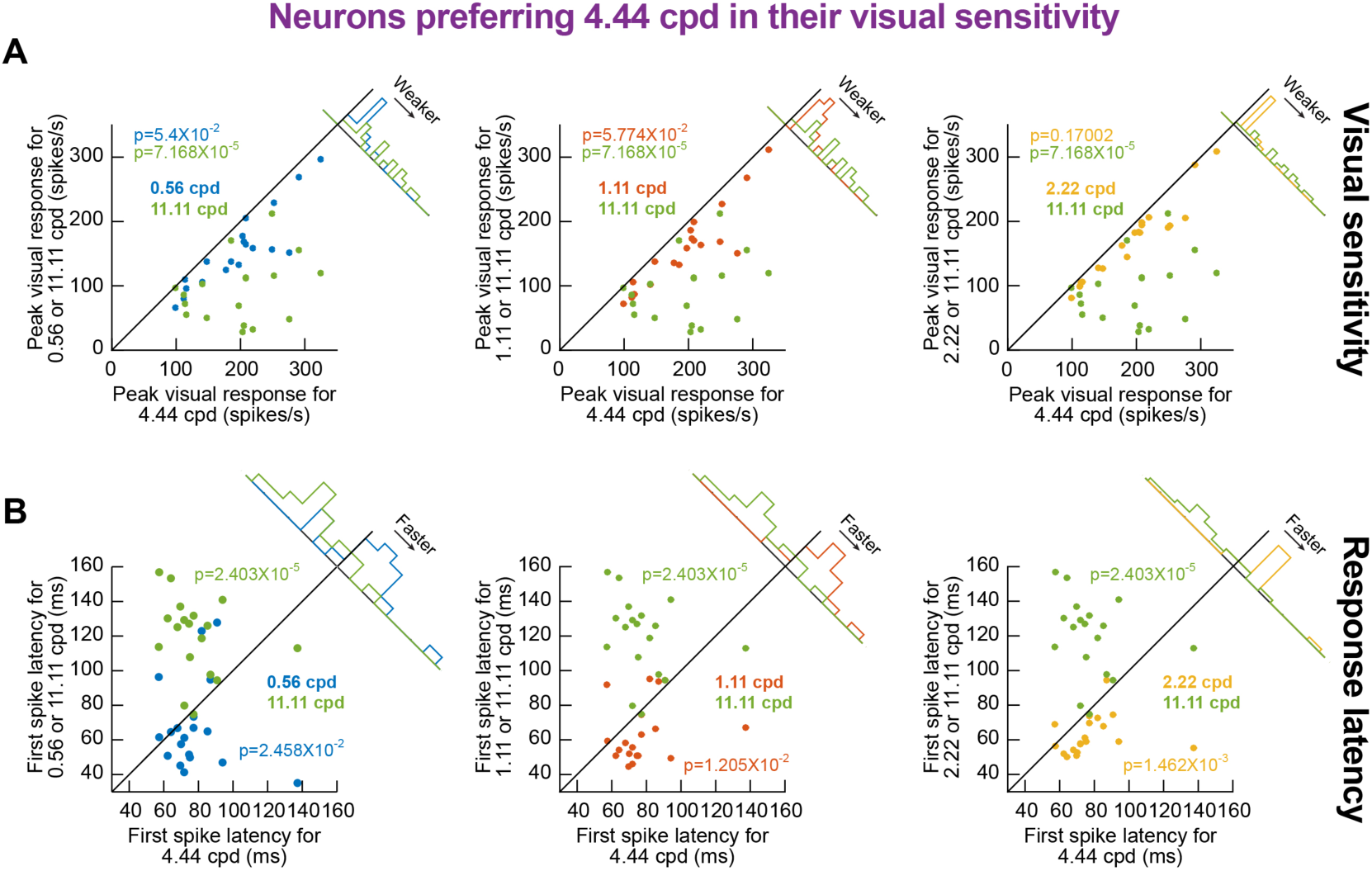
Rapid detection of low spatial frequencies in the macaque SC independent of neural sensitivity. (**A**) For neurons showing the highest visual responses to 4.44 cpd gratings (N=19), we plotted in each panel visual response strength for either higher or lower spatial frequencies (y-axis) against response strength for 4.44 cpd. For example, in the leftmost panel, the bluish dots show responses to 0.56 cpd versus responses to 4.44 cpd, and the greenish dots show responses to 11.11 cpd versus responses to 4.44 cpd. As expected, response strength was always highest for 4.44 cpd. P-values are indicated in each panel, reflecting a comparison between either the higher or lower spatial frequency (color-coded according to the legend) to 4.44 cpd using a Ranksum test. (**B**) We measured first-spike latency for 4.44 cpd gratings (x-axis) and related it to first-spike latency for either lower or higher spatial frequencies (y-axis). Even though 4.44 cpd gratings always evoked the strongest visual response (**A**), first-spike latency for 4.44 cpd gratings was either longer or shorter than the latency for other gratings. Moreover, whether first-spike latency for the preferred spatial frequency (4.44 cpd) was longer or shorter than in other spatial frequencies simply depended on the rank-ordering of spike timing observed in Fig. 1. Thus, coarse-to-fine visual sensation by the SC is independent of response strength.

Thus, in terms of visual sensitivity (Fig. 2A), all spatial frequencies other than 4.44 cpd expectedly elicited weaker neural responses than 4.44 cpd, since all the neurons selected in this analysis preferred 4.44 cpd by definition. However, despite such preference, visual response latency (Fig. 2B) in the same neurons was either significantly shorter or longer than the latency observed for 4.44 cpd (p-values for statistical tests are shown in the figure), and following a very simple rule: for 0.56, 1.11, and 2.22 cpd spatial frequencies, response latencies were shorter than for 4.44 cpd, whereas response latencies were longer for 11.11 cpd. Again, for all these spatial frequencies, response sensitivity was weaker than for 4.44 cpd. Thus, faster SC detection of low spatial frequencies occurs independently of visual sensitivity to a given spatial frequency.

This observation also persisted when we considered neurons preferring other spatial frequencies. In Fig. 3, we plotted visually-evoked responses for different spatial frequencies, but after separating neurons in each panel according to their preferred spatial frequency in terms of response sensitivity. The leftmost panel shows neurons responding the strongest for 0.56 cpd, and the rightmost panel shows neurons responding the strongest for 4.44 cpd, and so on for the other panels. Yet, and as can be seen from the arrows indicating the times of peak visual responses for each spatial frequency, the lowest two spatial frequencies always evoked the fastest responses followed by a systematic increase in response latency with increasing spatial frequency; again, this happened regardless of neural preference for a given spatial frequency in terms of sensitivity.

**Figure 3.**
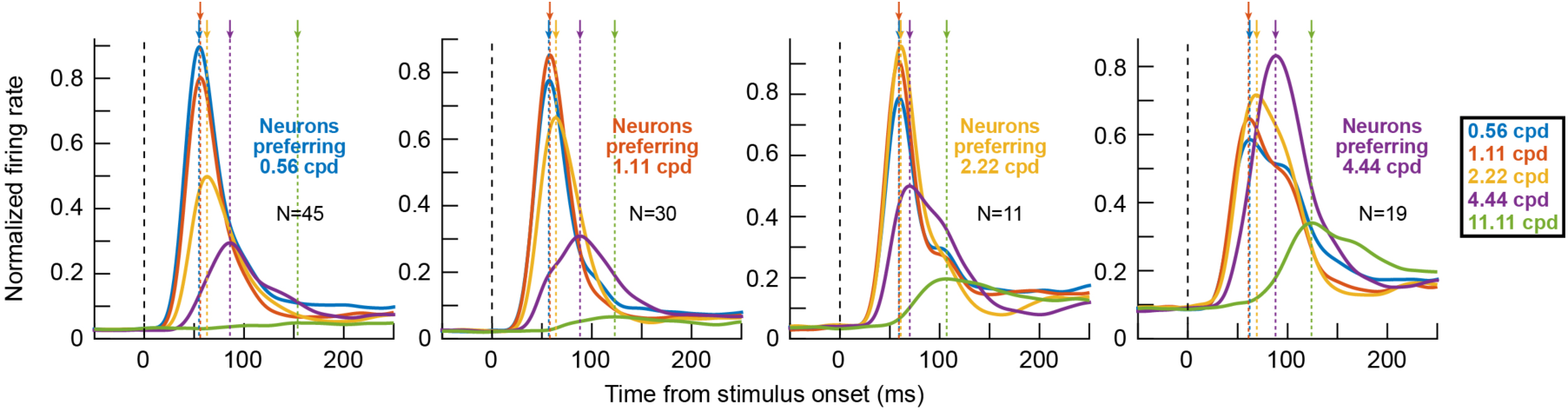
Dissociation between SC response strength and latency as a function of spatial frequency. The effect in Fig. 2 persisted for neurons preferring other spatial frequencies. In each panel, we took only neurons preferring one spatial frequency in terms of their visual sensitivity. As expected, gratings of non-preferred spatial frequencies emitted weaker visual responses than the preferred spatial frequency in each panel. However, regardless of visual sensitivity to a given spatial frequency, the rank ordering of visual burst times as a function of spatial frequency was similar across all panels (indicated by the downward arrows highlighting the time of peak visual response for each spatial frequency). For example, the responses to 2.22 and 4.44 cpd gratings always came later than the responses to 0.56 and 1.11 cpd gratings regardless of which spatial frequency the neurons preferred. Note that we did not have enough neurons preferring 11.11 cpd to include in this analysis (see Figs. 5–6). The numbers of neurons contributing to each panel are indicated in the figure.

We next analyzed the population-level properties of SC visual response latencies by plotting cumulative histograms of first-spike latency after grating onset (Materials and Methods). Across the population, the lowest two spatial frequencies (0.56 and 1.11 cpd) consistently evoked the shortest visual response latencies followed by a monotonic increase with increasing spatial frequency (Fig. 4A, B, including statistical tests).

**Figure 4.**
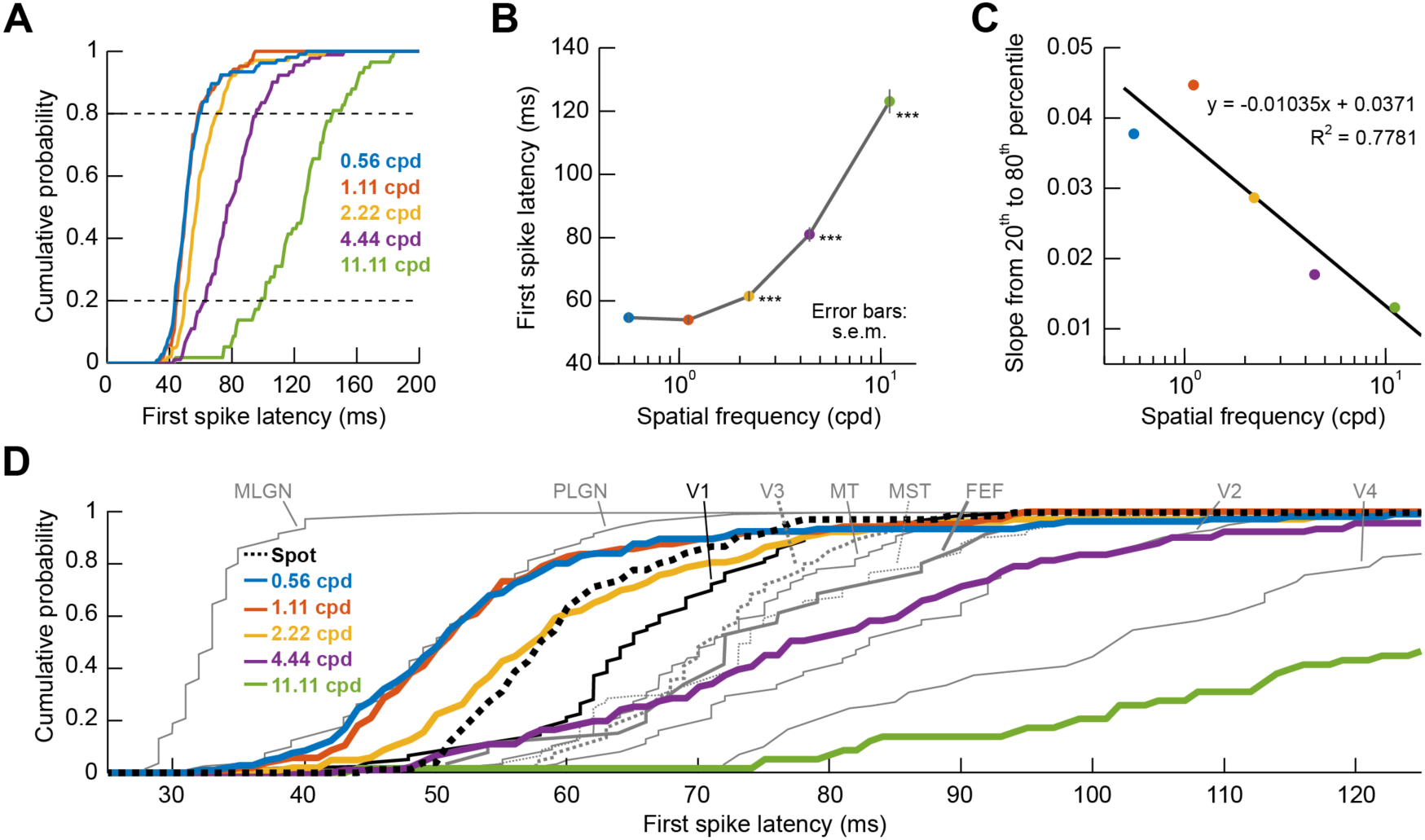
Early visual sensation by the macaque SC. (**A**) Cumulative histograms of first-spike latency (Materials and Methods) in our recorded neurons, separated by spatial frequency. For each neuron, we measured the average first-spike latency of the evoked visual response after a given spatial frequency grating was presented on multiple trials. We then repeated the measurement for other spatial frequencies. The evoked response consistently came earlier for low spatial frequencies than for high spatial frequencies. (**B**) The rank-ordering of spatial frequencies in **A** is also seen when plotting the mean first-spike latency across all neurons as a function of spatial frequency. Low spatial frequencies evoked a visual response earlier than high spatial frequencies. Error bars denote s.e.m. across neurons, and the asterisks indicate p<0.001 when comparing first-spike latency for 0.56 cpd to that in each of the other spatial frequencies. (**C**) Variability of first-spike latency was higher for higher spatial frequencies. We plotted the slope of the cumulative histograms in **A** between the 20^th^ and 80^th^ data percentiles as a function of spatial frequency. This slope progressively decreased, suggesting progressive increase in first-spike latency variability across neurons with higher spatial frequencies. (**D**) Our data from **A** plotted along with data from multiple visual areas in gray; copied with permission from (Schmolesky et al., 1998). SC visual responses for low spatial frequencies were among the earliest responses in the visual system, but it has to be noted that the Schmolesky et al. data were from anesthetized animals (i.e. exhibiting slightly delayed visual responses when compared to awake ones); also see (Huang and Paradiso, 2008) for example awake-monkey V1 latencies. The thick dotted curve shows the distribution of first-spike latencies in our neurons when a small spot of light was presented instead of a grating (i.e. a stimulus with broad-band spatial frequency). Also see Figure Supplements 1 and 2.

Moreover, this increase was accompanied by increased latency variability across neurons (Fig. 4C), and it persisted for either purely visual or visual-motor SC neurons (Fig. 4-Figure Supplement 1). This effect was also independent of differences in response latency between upper and lower visual field SC representations (Hafed and Chen, 2016), because the impact of spatial frequency on response latency still occurred even after we separated neurons as either representing the upper or lower visual fields (Fig. 4-Figure Supplement 2).

Interestingly, we found that SC visual response latencies were among the earliest in the whole visual system based on the published literature, and particularly for low spatial frequency stimuli. For example, Schmolesky and colleagues (Schmolesky et al., 1998) characterized visual response latencies in lateral geniculate nucleus (LGN) and a variety of cortical areas. These authors used large spots or bars filling each neuron's RF, resulting in a broad spectrum of low and high spatial frequencies in their stimuli. When we plotted our observed SC visual response latencies along with these authors’ results as a reference (Fig. 4D), we found that the SC consistently exhibited very early responses (for example, compare our 2.22 cpd latencies to those in V1 from their measurements). Moreover, even when we measured SC visual response latencies using small bright spots, that is, still activating a broad spectrum of spatial frequencies, the SC still exhibited early latencies (for example, compare the dashed and dotted black lines). Even though the Schmolesky data were collected using anesthetized animals, in which response latencies are delayed relative to the awake condition (Vaiceliunaite et al., 2013), our results nonetheless indicate that SC visual responses are indeed among the most rapidly evolving responses in the entire visual system (Fig. 4D; also see Huang and Paradiso, 2008 for example awake-monkey V1 latencies), and this effect is magnified even further when low spatial frequencies are present in the stimuli.

### Over-representation of low spatial frequencies in SC neural sensitivity

Besides rapidly detecting low spatial frequencies, being able to efficiently guide behavior implies that the SC's pattern analysis machinery might also be more sensitive to such low spatial frequencies, at the population level, and not just be faster in responding to them. Indeed, when we plotted all tuning curves as done in Fig. 1C (top), we found primarily low-pass characteristics in the population even at near-foveal eccentricities. Specifically, Fig. 5A shows sensitivity tuning curves of 3 example neurons from different retinotopic eccentricities, and Fig. 5B, C summarizes the population results. The range of preferred spatial frequencies was expectedly higher at near-foveal eccentricities than at extra-foveal ones (Fig. 5C) (Hafed and Chen, 2016), but the overall population curves were primarily low-pass (black curves in Fig. 5B). This suggests that the SC over-represents low spatial frequencies in terms of visual response sensitivity in addition to its boosting of such spatial frequencies in terms of response latency.

**Figure 5.**
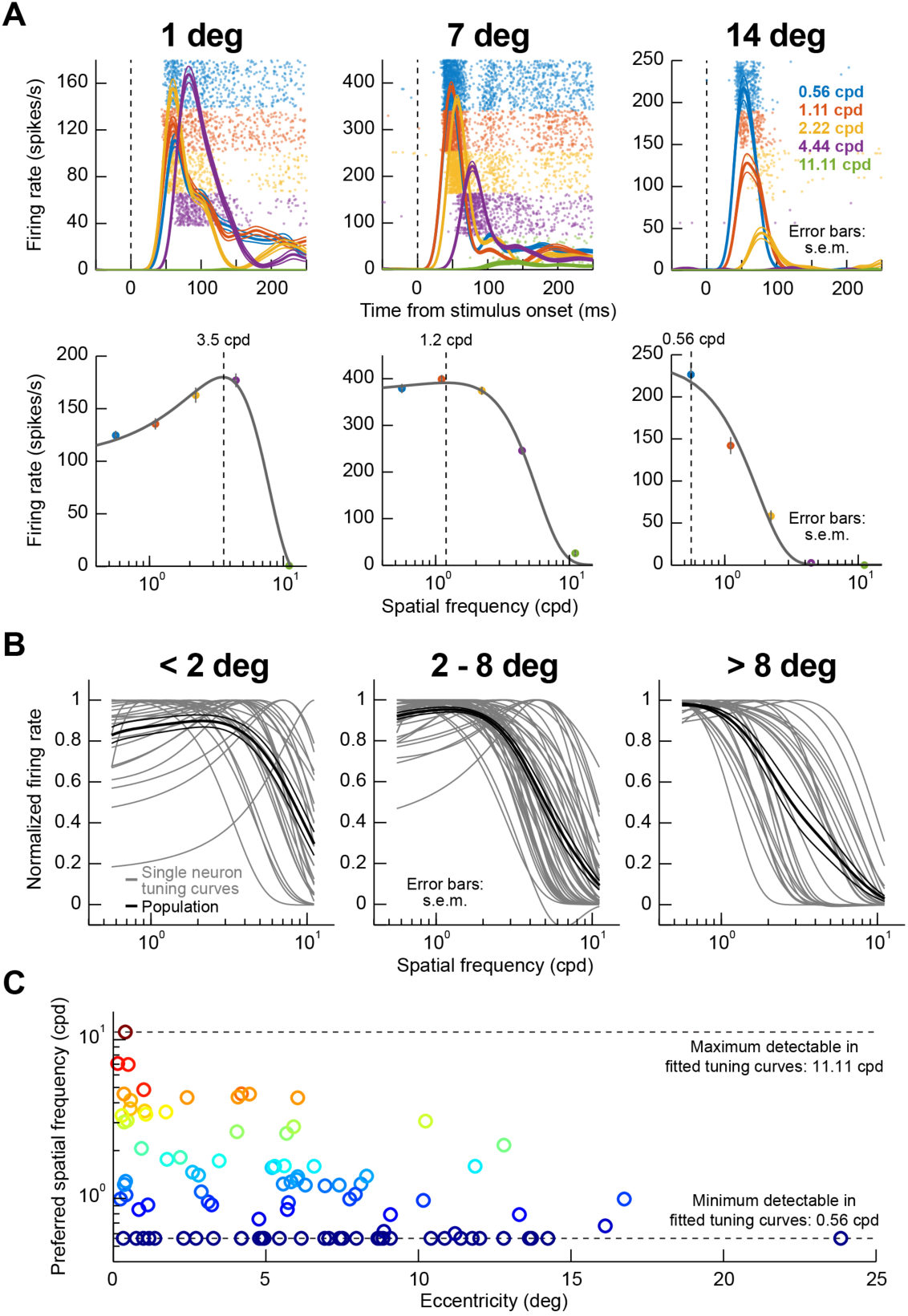
Predominantly low-pass spatial frequency preference in terms of neural sensitivity. (**A**) Example visual responses of 3 SC neurons preferring different retinotopic eccentricities (1, 7, or 14 deg). Each panel in the top row plots firing rate as a function of time from stimulus onset for gratings presented within each neuron's visual RF; color codes indicate the spatial frequency of the presented stimulus. Raster plots in the background show times of action potentials across trials. Firing rate curves show mean and s.e.m. as thick and thin lines, respectively. The near-foveal neuron (1 deg) preferred higher spatial frequencies than the more eccentric neurons, as evidenced by the higher responses for 4.44 cpd gratings than for lower spatial frequencies in this neuron. The bottom panels show spatial frequency tuning curves for the same neurons. The dashed vertical line in each panel indicates the preferred spatial frequency of each neuron based on the tuning curves. (**B**) Tuning curves from all neurons in our population, grouped into 3 different eccentricity bins. Thin curves show individual neuron tuning curves, and thick black curves show the mean tuning curve within a given panel, along with s.e.m. error bars across neurons as thin black lines. Regardless of eccentricity, population tuning curves showed primarily low-pass characteristics (thick black curves), and this effect got stronger the more eccentric the neurons were (compare panels). (**C**) Preferred spatial frequency as a function of neuronal preferred eccentricity. Near-foveal neurons had a broad range of preferred spatial frequencies, but there was still low spatial frequency preference at these eccentricities. Preferred spatial frequency was selected in this figure as the peak in fitted tuning curves, like those shown in **A**. Thus, for extremely low- or high-pass neurons, the preferred spatial frequency indicated in this analysis was only an estimate that was cut-off by the end of the fitted curves constrained by our sampled spatial frequencies (dashed horizontal lines).

We explored the over-representation of low spatial frequencies further by first counting the number of neurons responding the most for 0.56 cpd spatial frequencies as opposed to other spatial frequencies. These neurons accounted for 42% of our population, and no other single spatial frequency recruited as many neurons (Fig. 6A). Interestingly, this over-representation of low spatial frequencies became even more obvious when assessing local population activity reflected in field potentials (LFP's), which we recorded simultaneously around our electrode tips along with the isolated neurons (Materials and Methods). We measured either the evoked (Fig. 6B) or sustained (Fig. 6C) local population activity after grating onset (Materials and Methods), and the great majority of our electrode locations (64% for the transient response and 77% for the sustained response), whether in near-foveal or extra-foveal locations, picked up the strongest responses for the lowest spatial frequency that we presented (Fig. 6B, C). This effect can also be better appreciated when inspecting raw LFP traces from the same 3 example electrode penetrations from which the 3 example neurons of Fig. 5A were isolated (Fig. 6D). Even though the near-foveal neuron in Fig. 5A (leftmost panel) responded the most for the 4.44 cpd grating, the local population picked up by the LFP signal around the electrode in the same experiment still showed the strongest stimulus-evoked deflection (as well as sustained response) for 0.56 and 1.11 cpd gratings (Fig. 6D, leftmost panel). In other words, at the population level, even near-foveal SC eccentricities over-represent low spatial frequencies. Similar effects were also observed for the other two example eccentricities in Fig. 6D. Therefore, the SC over-represents low spatial frequencies both in terms of neural sensitivity (Figs. 5–6) as well as neural response latency (Figs. 1–4).

**Figure 6.**
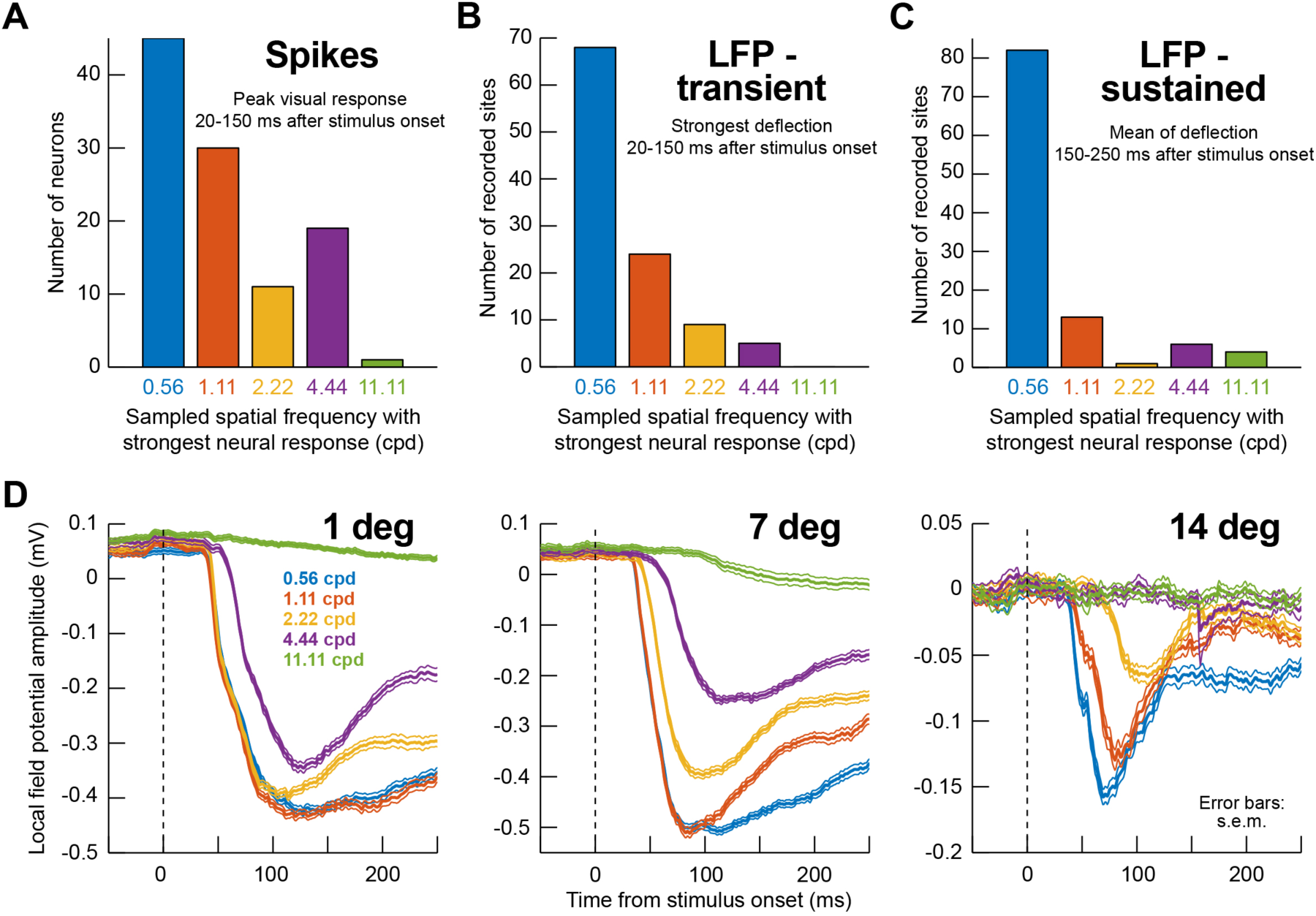
Low-pass spatial frequency filtering characteristics of the macaque SC. (**A**) Distribution of preferred spatial frequencies in our population of recorded SC neurons. In this analysis, we binned neurons according to the presented spatial frequency grating that elicited maximal neuronal response. More neurons were driven the strongest by the lowest spatial frequency (0.56 cpd) than by any other higher spatial frequency. (**B**) We performed a similar analysis but on the transient evoked local field potential (LFP) response (Materials and Methods; also see **D** for example evoked LFP responses, which are negative going). The number of electrode penetrations that showed maximal response for 0.56 cpd was even higher than for the isolated neurons in **A**. (**C**) This effect was even stronger in the sustained LFP response starting after 150 ms from stimulus onset. Thus, at the population level reflected by LFP signals, the SC is primarily tuned to low spatial frequencies. (**D**) Stimulus-evoked LFP responses from the same electrode penetrations in which the example neurons of Fig. 5A were isolated and recorded. In the LFP, all 3 electrode tracks, regardless of eccentricity, showed a preference for low spatial frequencies (stronger negative deflections), even in the near-foveal SC region where the neuron preferring 3.5 cpd in Fig. 5A was isolated. This means that the SC over-represents low spatial frequencies in neural sensitivity. Error bars denote s.e.m.

### Faster scanning of low spatial frequencies with saccadic eye movements

Finally, we related SC visual response latency and sensitivity to behavior. In completely separate behavioral sessions, we trained our monkeys to generate visually guided saccades to gratings of different spatial frequencies (Chen and Hafed, 2017). As expected, saccadic reaction time (RT) increased with increasing spatial frequency in each animal (Fig. 7C, F; black curves) (Chen and Hafed, 2017; Ludwig et al., 2004), and our interest here was in how such RT increases were correlated with SC neural visual response properties (even when these neural visual responses were recorded in the complete absence of saccadic orienting to the gratings, and in different experimental sessions). We thus plotted SC visual response strength (Fig. 7A, D) and latency (fig. 7B, E) for each monkey's neurons individually (again, the monkeys never made target-directed saccades to the gratings during the recordings). We found that a simple linear combination (Materials and Methods) of visual response strength and latency in a given monkey allowed predicting this monkey's behavior (Fig. 7C, F; green curves) remarkably well (r^2^: 99.98% for monkey N and 91.91% for monkey P). Moreover, the model prediction (i.e. r^2^) was better than when using either response strength alone (99.27% for monkey N and 90.86% for monkey P) or response latency alone (99.5% for monkey N and 88.34% for monkey P). Thus, SC visual response properties are strongly related to how efficiently saccades can be made to different spatial frequencies, even when neural and behavioral correlations are performed from completely different experimental sessions.

**Figure 7.**
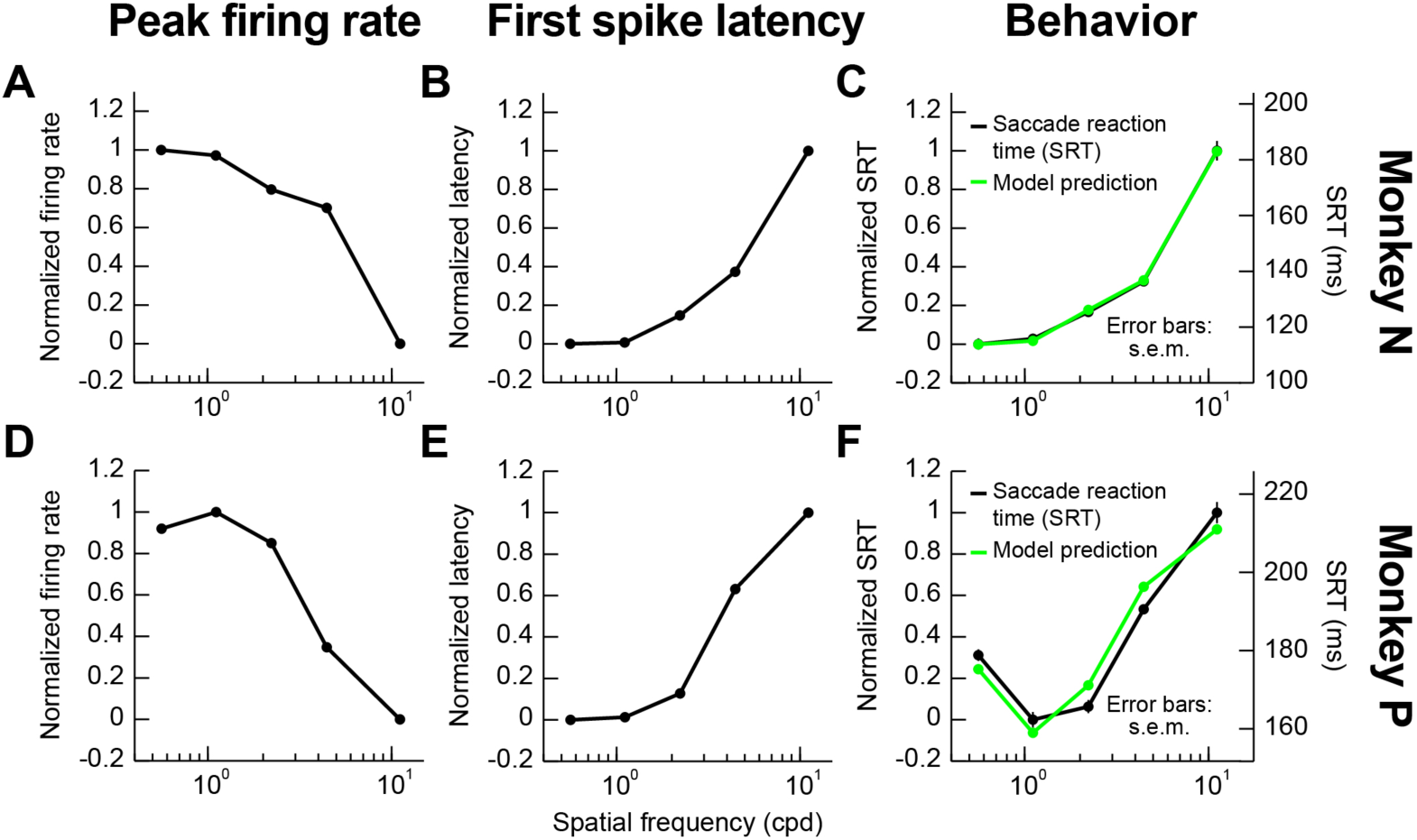
Linking visual response latency and strength to saccade behavior. (**A**) For monkey N, we plotted the average response strength as a function of spatial frequency for all neurons covering an eccentricity similar to an eccentricity used in separate behavioral sessions requiring a saccade to the gratings (Materials and Methods). Firing rates were normalized in the range of 0-1. Low spatial frequencies were associated with higher responses, as shown in Figs. 5–6. (**B**) We performed a similar analysis for neural response latency from the same neurons; this time, low spatial frequencies were associated with more rapid neural responses, as shown in Figs. 1–4. (**C**) The black curve shows the monkey's saccade reaction time (RT) from completely different behavioral sessions, and the green curve shows a linear combination of the neural curves in **A**, **B**. As can be seen, behavioral performance matched neural performance well. (**D-F**) Similar analyses for monkey P. Error bars are defined in the figure where appropriate. For monkey N, model parameters a, b, and c from equation 2 in Materials and Methods were −0.458, 0.537, and 0.465, respectively. For monkey P, they were −1.959, −1.058, and 1.985, respectively.

We then wanted to relate our results above to human performance. We performed human behavioral experiments testing the predictions of our SC-based observations. We ran a visual search task exercising different spatial frequencies in active gaze behavior, but with highly visible stimuli even at high spatial frequencies. Subjects had to freely search for a grating with an oddball orientation from among many other ones having the same spatial frequency but a slightly different orientation from the oddball stimulus (Fig. 8A; Materials and Methods). The task was demanding enough that subjects had to generate many saccades to search for the oddball target, and example scan paths of these saccades are shown in green in Fig. 8A. We found that inter-saccadic intervals increased in duration when the search array consisted of high spatial frequencies as opposed to low spatial frequencies (Fig. 8B), consistent with our neural and behavioral results above. Importantly, this effect was not due to a speed-accuracy tradeoff, in which it may have been the case that faster inter-saccadic intervals were associated with worse task performance. Instead, Fig. 8C demonstrates that target detection performance was almost constant (and at high levels) for the spatial frequencies (for example, between ~1 and ~4 cpd) in which inter-saccadic intervals showed the biggest changes in duration. Moreover, for all but the very last 1-2 inter-saccadic intervals in Fig. 8B, the oddball target was not yet identified by the subjects, so the shorter intervals for low spatial frequencies were not because oddball targets were already identified or recognized. Therefore, even in natural searching gaze behavior with a stimulus that is less prone to visual masking as in (White et al., 2008), the effects of spatial frequencies on inter-saccadic intervals are consistent with a role of the SC (and other brain areas) in facilitating and over-representing low spatial frequencies prevalent in natural environments.

**Figure 8.**
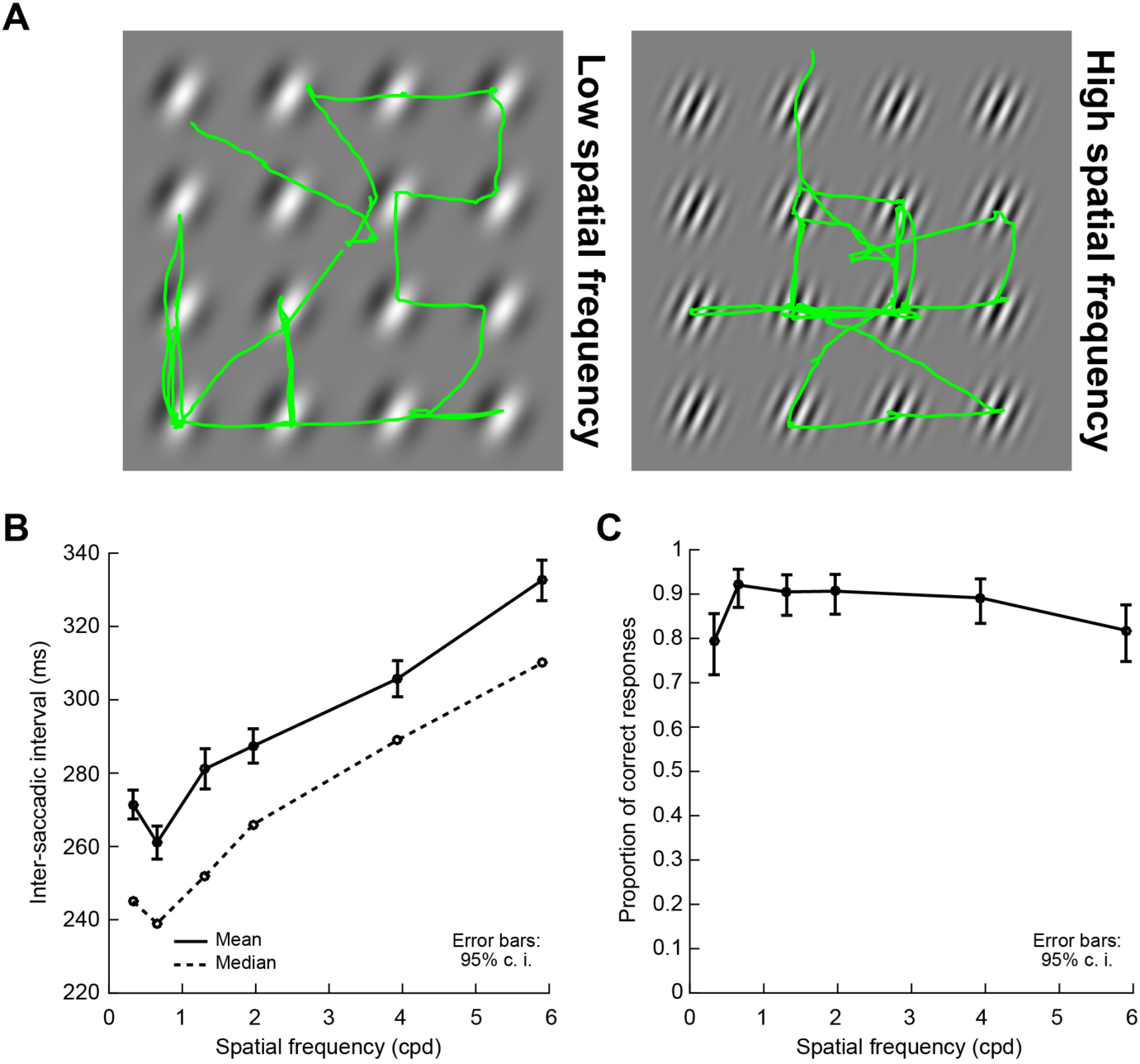
Influence of spatial frequency on inter-saccadic intervals during human visual search behavior. (**A**) Example visual search arrays and eye movement scan paths (green) superimposed on them. The left panel shows a search array of targets with a low spatial frequency, and the right panel shows a search array of targets with a higher spatial frequency. In all cases, subjects searched for an oddball orientation in the array, which was slightly different from the orientation that was present in all other gratings; the task was made difficult enough to require many scanning saccades of the array before the correct oddball stimulus was identified. (**B**) Mean (solid) and median (dashed) inter-saccadic intervals during target search as a function of the spatial frequency of the targets in the search array. Error bars denote 95% confidence intervals. Inter-saccadic intervals progressively increased with higher spatial frequencies, as in our earlier SC neural and behavioral results. (**C**) Proportion of correct oddball identifications (Materials and Methods) as a function of spatial frequency in the target search array. Subjects faithfully searched the array until they could correctly identify the target on the great majority of trials, meaning that the changes in inter-saccadic intervals in **B** were not due to potential speed-accuracy tradeoffs during search. Error bars denote 95% confidence intervals.

## Discussion

We found preferential representation of low spatial frequencies in the SC in terms of both response latency and response strength. We believe that these results place the SC in an ideal position to facilitate orienting behavior in natural environments, which are dominated by low image spatial frequencies (Ruderman and Bialek, 1994; Tolhurst et al., 1992). Consistent with this, White and colleagues recently found that saccadic RT's are significantly faster in natural scenes, after ensuring matched stimulus visibility (White et al., 2008). We also performed a visual search task with high target visibility (Fig. 8), and we found strong dependence of saccadic timing on spatial frequency, consistent with our SC results. In this regard, our analysis of monkey saccadic RT's in Fig. 7 is particularly intriguing, because it further suggests that the SC may indeed be instrumental in facilitating these human observations. Specifically, we were able to account for each animal's RT patterns with high fidelity based solely on SC visual responses collected from different experimental sessions and during passive fixation. We believe that this result makes sense in hindsight. If one were to expect any visual brain areas to be optimized for natural scene statistics dominated by low spatial frequencies, then it should be those areas that have privileged access to the motor output, like the SC. This is imperative given that we are active observers and would normally orient our sensory apparati very frequently under natural scene scenarios.

Our results are also interesting because they highlight the importance of the primate SC's visual functions. Historically, this structure was studied heavily from the perspective of motor control (Gandhi and Katnani, 2011), with more recent interest focusing on cognitive processes related to attention, decision making, and target selection (Basso and May, 2017; Krauzlis et al., 2004; Krauzlis et al., 2013). However, the SC is also a visual structure, and it is the primary visual structure in lower animals. Indeed, recent results have examined visual properties of the primate SC, for example to color stimuli, more closely (Hall and Colby, 2016; Herman and Krauzlis, 2017; White et al., 2009). We particularly think that our results may provide a possible mechanism for allowing the SC to preferentially process face-like stimuli, as was also recently observed (Nguyen et al., 2014), and to mediate known preferential orienting patterns to such stimuli (Johnson, 2005; Ro et al., 2007; Tomalski et al., 2009). Moreover, the SC's visual functions could be important in supporting an alternative visual pathway during blindsight (Weiskrantz et al., 1974). In this condition, patients with V1 loss lose conscious perception but nonetheless exhibit residual sensory, cognitive, and motor capabilities that may be mediated by pathways bypassing V1 (for example, direct retinal projections to the SC). Indeed, some primate models of blindsight point to the importance of the SC in guiding saccades under this condition (Kato et al., 2011; Takaura et al., 2011; Yoshida et al., 2008). However, what is perhaps more intriguing to us is that studies of spatial frequency sensitivity of human blindsight patients show spatial frequency cutoffs near ~4 cpd (Sahraie et al., 2010; Sahraie et al., 2002; Trevethan and Sahraie, 2003), which is remarkably similar to the capabilities of SC neurons that we have observed in our data (Fig. 5B). Thus, our results could help clarify the conditions under which the SC may be particularly important for blindsight performance when compared to other visual pathways that also bypass V1 (for example, geniculo-cortical pathways).

Related to spatial frequency cutoffs, we found primarily low-pass characteristics of SC neurons (Figs. 5–6). Of course, at near-foveal eccentricities, where higher resolution vision dominates, band-pass tuning curves were observed, but the overwhelming population result was low-pass in nature. In the SC of anesthetized marmosets, band-pass tuning was suggested in (Tailby et al., 2012), although it is not clear how this may have depended on eccentricity in that study. These authors also observed low-pass tuning in some of their own neurons. In any case, our observations of primarily low-pass tuning curves is reminiscent of LGN spatial frequency tuning curves (Kaplan and Shapley, 1982) rather than V1 ones, which tend to be band-pass (De Valois et al., 1982)

Potential similarities between the SC and LGN may also extend to visual response latencies. In Fig. 4D, we plotted, just as a reference, observations of visual response latencies from (Schmolesky et al., 1998), and we found that for low spatial frequencies, SC visual responses may arrive earlier than in V1. Of course, the fact that monkeys were anesthetized in that earlier study does not allow for a proper quantitative comparison, because anesthesia delays visual responses (Vaiceliunaite et al., 2013). However, even comparison of our results to awake macaque V1 latencies in (Huang and Paradiso, 2008) would suggest that the SC is at least as fast as V1, if not slightly faster, in detecting visual stimuli. This is consistent with the SC receiving direct retinal projections (Pollack and Hickey, 1979) and also with the fact that eye movements, including microsaccades, can be reflexively altered by visual stimuli with latencies much earlier than the latencies of most cortical visual areas (Buonocore et al., 2017; Edelman and Keller, 1996; Hafed et al., 2015; Hafed and Ignashchenkova, 2013; Tian et al., 2016).

In all, we believe that our results demonstrate that spatial vision capabilities of the primate SC are specifically organized to facilitate exploring natural scenes with rapid gaze shifts.

## Materials and Methods

Monkey experiments were approved by regional governmental offices in Tuebingen. For the human experiments, ethics committees at Tuebingen University reviewed and approved our protocols. All human subjects provided written informed consent in accordance with the Declaration of Helsinki.

## Animal preparation

Monkeys P and N (male, *Macaca mulatta*, aged 7 years) were prepared for behavior and superior colliculus (SC) recordings earlier (Chen and Hafed, 2013; Chen et al., 2015; Chen and Hafed, 2017). Briefly, we placed a recording chamber centered on the midline, and we angled it to point towards a stereotaxically defined point 1 mm posterior of and 15 mm above the inter-aural line. The chamber angle was posterior of vertical (by 38 and 35 deg for monkeys P and N, respectively).

## Monkey recording task

The monkeys performed a pure fixation task while we recorded the activity of visually-responsive SC neurons, as described in detail before (Chen and Hafed, 2017; Chen et al., 2015). Briefly, in each trial, we displayed a white fixation spot (8.5×8.5 min arc) over a gray background. Fixation spot and background luminance were described earlier (Chen and Hafed, 2013). After an initial fixation interval (400-550 ms), the fixation spot transiently dimmed for ~50 ms, which reset microsaccadic rhythms (Hafed and Ignashchenkova, 2013; Tian et al., 2016) and also attracted attention to the spot rather than to the response field (RF) stimulus. After an additional 110-320 ms, a stationary, vertical Gabor patch with 80% relative contrast (defined as Lmax-Lmin/Lmax+Lmin) appeared for 300 ms within the neuron's RF. The RF was estimated earlier in the session using standard saccade tasks (Chen et al., 2015; Hafed and Chen, 2016), and the Gabor patch size was chosen to fill as much of the RF as possible. The spatial frequency of the patch, in cycles/deg (cpd), was varied randomly across trials (from among 0.56, 1.11, 2.22, 4.44, and 11.11 cpd). Grating phase was randomized from trial to trial, and the monkey was rewarded only for maintaining fixation; no orienting to the grating or any other behavioral response was required. We used only vertical gratings, but we confirmed that they elicit robust responses in the SC. In pilot data, we also confirmed that any potential orientation tuning in the SC was broad and included robust responses to vertical gratings (Chen et al., 2015; Marrocco and Li, 1977; Schiller and Koerner, 1971).

We recorded from 115 neurons (monkey N: 60; monkey P: 55) with preferred eccentricities up to 24 deg. We excluded trials with microsaccades occurring within +/- 100 ms from stimulus onset because such occurrence can alter neural activity. In fact, the trials with microsaccades near stimulus onset were analyzed recently, from the same set of neurons, to explore spatial-frequency dependence of saccadic suppression in the SC (Chen and Hafed, 2017). Our focus here was to only analyze baseline visual activity and not activity modulated due to the presentation of peri-movement stimuli. We excluded 9 neurons from further analyses because they did not have >25 repetitions per tested spatial frequency after excluding the microsaccade trials. This number was our chosen threshold for the minimum number of observations in order to have sufficient confidence in our interpretations of the results. For the remaining 106 neurons that were included in the analyses, we collected >295 trials per neuron (average: 935 +/- 271 s.d.).

## Monkey saccade reaction time task

In completely different purely behavioral sessions, we ran our monkeys on a simple saccade reaction time task, which we recently described in detail (Chen and Hafed, 2017). Briefly, the monkeys fixated, and a Gabor patch of 2 deg diameter could appear at 3.5 deg eccentricity either to the right or left of fixation. The patch was otherwise identical to that used in the recording task described above, and the fixation spot disappeared simultaneously with patch appearance in order to cue the monkeys to generate a targeting saccade towards the patch. We measured reaction time (RT) and correlated it with SC visual responses collected from completely different sessions and critically not involving a saccadic response at all (i.e. the recording task above). We analyzed 2,522 trials from Monkey N and 3,392 trials from Monkey P. As with the neural data above, we only analyzed trials without any microsaccades within 100 ms before or after Gabor patch onset, to avoid peri-movement effects on RT that were described in detail elsewhere from the same experimental sessions (Chen and Hafed, 2017).

## Human visual scanning task

Subjects sat in a dark room facing a computer display (41 pixels per deg; 85 Hz), and head fixation was achieved through a custom-made chin/forehead rest (Hafed, 2013). We collected data from 8 subjects (3 females and 5 males; 5 subjects were authors of the study).

Each trial started with an initial fixation spot presented at display center. After ~1030 ms of steady fixation, a search array consisting of 4×4 Gabor patches appeared. Each patch was 6.1 deg in diameter, and all 16 patches were distributed evenly in a grid layout across the display. Grating contrast was set to maximum (100%), and all patches had the same spatial frequency within a given trial. Spatial frequency was altered randomly across trials from among 6 possible values (0.33, 0.66, 1.31, 1.97, 3.93, and 5.9 cpd). Moreover, all but one patch had the same orientation within a given trial (picked randomly across trials from all possible orientations with a resolution of 1 deg). The odd patch was tilted by 7 deg either clockwise or counter-clockwise from the orientation of all other patches, and the subjects’ task was to search for the oddly oriented patch and indicate whether it was tilted to the right or left from all other patches. The task was very difficult to perform during fixation, and therefore required prolonged scanning of the entire grid array of patches with many saccades until the odd patch was found and correctly discriminated. This allowed us to obtain sufficient search performance data, with many inter-saccadic intervals that were the focus of our analysis (i.e. our goal was to investigate how inter-saccadic intervals were affected by spatial frequency). We collected 180 trials per subject (i.e. 30 trials per spatial frequency), but each trial had many more inter-saccadic intervals that could be analyzed (as detailed below).

## Neuron classification

We used similar neuron classification criteria to those used in our recent studies (Chen et al., 2015; Hafed and Chen, 2016). Briefly, a neuron was labeled as visual if its activity 0-200 ms after target onset in a delayed saccade task (Hafed and Chen, 2016; Hafed and Krauzlis, 2008) was higher than activity 0-200 ms before target onset (p<0.05, paired t-test). The neuron was labeled as visual-motor if its pre-saccadic activity (-50-0 ms from saccade onset) was also elevated in the delayed saccade task relative to an earlier fixation interval (100-175 ms before saccade onset) (Li and Basso, 2008). Our results (e.g. spatial frequency rank ordering of response latencies) were similar for either visual or visual-motor neurons (except for small quantitative differences in visual response latency). As a result, we combined neuron types in analyses unless otherwise explicitly stated.

## Eye movement analyses

We measured eye movements in monkeys using scleral search coils (Fuchs and Robinson, 1966; Judge et al., 1980), and we used a video-based eye tracker (EyeLink 1000, SR Research, Canada) for humans (Hafed, 2013). We detected saccades and microsaccades using velocity and acceleration criteria detailed elsewhere (Buonocore et al., 2017; Chen and Hafed, 2013; Hafed et al., 2009; Hafed and Ignashchenkova, 2013).

For the monkey recordings, we detected microsaccades in order to exclude trials with such movements occurring near stimulus onset (see above). For the monkey saccade reaction time task, we detected the targeting saccade after grating onset and measured its RT. We only considered trials in which there were no microsaccades within +/-100 ms from target onset, because microsaccades near target onset alter RT (Chen and Hafed, 2017; Hafed and Krauzlis, 2010), and because these trials with peri-microsaccadic stimuli were analyzed separately elsewhere (Chen and Hafed, 2017).

For the human scanning task, we measured inter-saccadic intervals during search. The inter-saccadic interval was defined as the time period between the offset of one saccade and the onset of the next. We only considered saccades occurring between search array onset and trial end (i.e. button press) when computing inter-saccadic intervals. Moreover, we only analyzed trials in which there were no blinks during the entire period from which we were collecting inter-saccadic intervals. Because trials were long until subjects found the odd patch, meaning that we had many inter-saccadic intervals within any trial, removal of blink trials did not reduce our data set dramatically; in the end, we had a total of 3,325-4,743 accepted inter-saccadic intervals per spatial frequency in our analyses (from a total of 145-182 accepted trials per spatial frequency).

## Firing rate analyses

We analyzed SC visual bursts by measuring peak firing rate 20-150 ms after stimulus onset (Chen and Hafed, 2017). We then obtained spatial frequency tuning curves by plotting peak visual response as a function of grating spatial frequency (Hafed and Chen, 2016). We performed a least squares fit of the measurements to the following difference-of-Gaussians function:

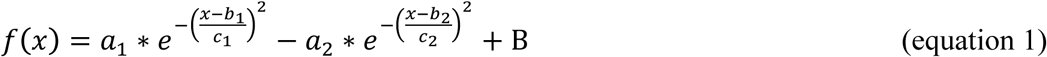

where f is firing rate, x is spatial frequency, a_1_ and a_2_ represent the amplitude of each Gaussian function, b_1_ and b_2_ represent the mean of each Gaussian function, c_1_ and c_2_ are the bandwidth of each Gaussian function, and B is the baseline firing rate (obtained from all trials as the mean firing rate in the interval 0-50 ms before Gabor patch onset). The goodness of fit was validated by computing the percentage of variance across stimuli accounted for by the model (Carandini et al., 1997). Only neurons that had >80% explained variance by the fit were included in summaries of tuning curve fits in Results (97 out of 106 neurons), but all neurons were included in all other analyses. We should also note here that the tuning curves from the same neurons in this study were presented earlier in brief format to provide support for the conclusions of another independent study out of our laboratory (Hafed and Chen, 2016); however, the conclusions and analyses shown in the present study are novel and were not described elsewhere before, whether by our laboratory or by other laboratories.

We estimated the preferred spatial frequency of each neuron as the spatial frequency within the sampled range of 0.56-11.11 cpd for which the fitted tuning curve from the above equation peaked. To combine different neurons’ tuning curves (e.g. Fig. 5B), we first normalized the peak of the tuning curve of each neuron to 1. We then combined neurons and obtained a mean curve across neurons along with s.e.m. estimates.

We estimated first-spike latency using Poisson spike train analysis (Legendy and Salcman, 1985). Most of our neurons had very little or no baseline activity, meaning that our estimate of first-spike latency using this method was very robust, and it gave us a sense of how quickly our neurons responded to the onset of a given stimulus.

## Local field potential analyses

We obtained local field potentials from wide-band neural signals using methods that we described recently (Chen and Hafed, 2017; Hafed and Chen, 2016). We then aligned LFP traces on Gabor patch onset, and we measured evoked responses in two ways. First, we measured the strongest deflection occurring in the interval 20-150 ms after stimulus onset, to obtain a measure that we called the transient LFP response. Second, we measured the mean deflection in the period 150-250 ms after stimulus onset, to obtain what we referred to as the sustained LFP response. Since the LFP evoked response is negative going, when we refer to a “peak” LFP response, we mean the most negative value of the measured signal.

## Predicting saccade reaction times from visual responses recorded on completely different sessions

Our approach was to ask whether RT from behavioral sessions can be related in a simple manner to visual response strength and first-spike latency from completely separate neural recording sessions in which no saccade to the patch was ever made. We used linear models of the form:

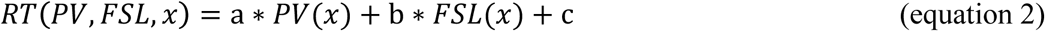

where x is spatial frequency, PV(x) is the average peak visual response of all included neurons for spatial frequency x; FSL(x) is the average first-spike latency of all included neurons for spatial frequency x; and a, b, c are model parameters. Since the behavioral RT's were experimentally obtained from horizontal targets at 3.5 deg eccentricity, we only included neurons with preferred RF locations centered within the range of 2-10 deg in eccentricity and +/-45 deg in direction from horizontal (i.e. 46 neurons). Moreover, we separated each monkey's neurons so that its own neural activity was used to predict its behavioral variability. Since we obtained similar conclusions when relating RT to either visual neurons alone or visual-motor neurons alone, we combined neuron types in the shown analyses to maximize the numbers of neurons used.

For modeling mean RT as a function of neural response properties, we normalized the range of RT values that we observed to the range from 0 to 1, with 0 corresponding to the shortest RT (e.g. that obtained from the lowest spatial frequency). We similarly normalized the range of peak visual response and first-spike latency. We then fit the best parameters to equation 2 above that matched the data. To test whether including either first-spike latency or peak visual response alone gave similar model fits to the case where both quantities were part of the model, we also ran the fitting with either parameter a or b in equation 2 pegged at 0.

## Author Contributions

C.-Y. C. and Z. M. H. performed the monkey experiments and analyzed the data. L. S., S. W., T. W., and Z. M. H. performed the human experiments. Z. M. H. wrote the paper.

## Acknowledgments

We were funded by the Werner Reichardt Centre for Integrative Neuroscience (CIN), an Excellence Cluster (EXC307) funded by the Deutsche Forschungsgemeinschaft (DFG). We were also supported by the Hertie Institute for Clinical Brain Research.

We declare no competing interests.

## Supplementary Figures

**Figure 4-Figure Supplement 1.**
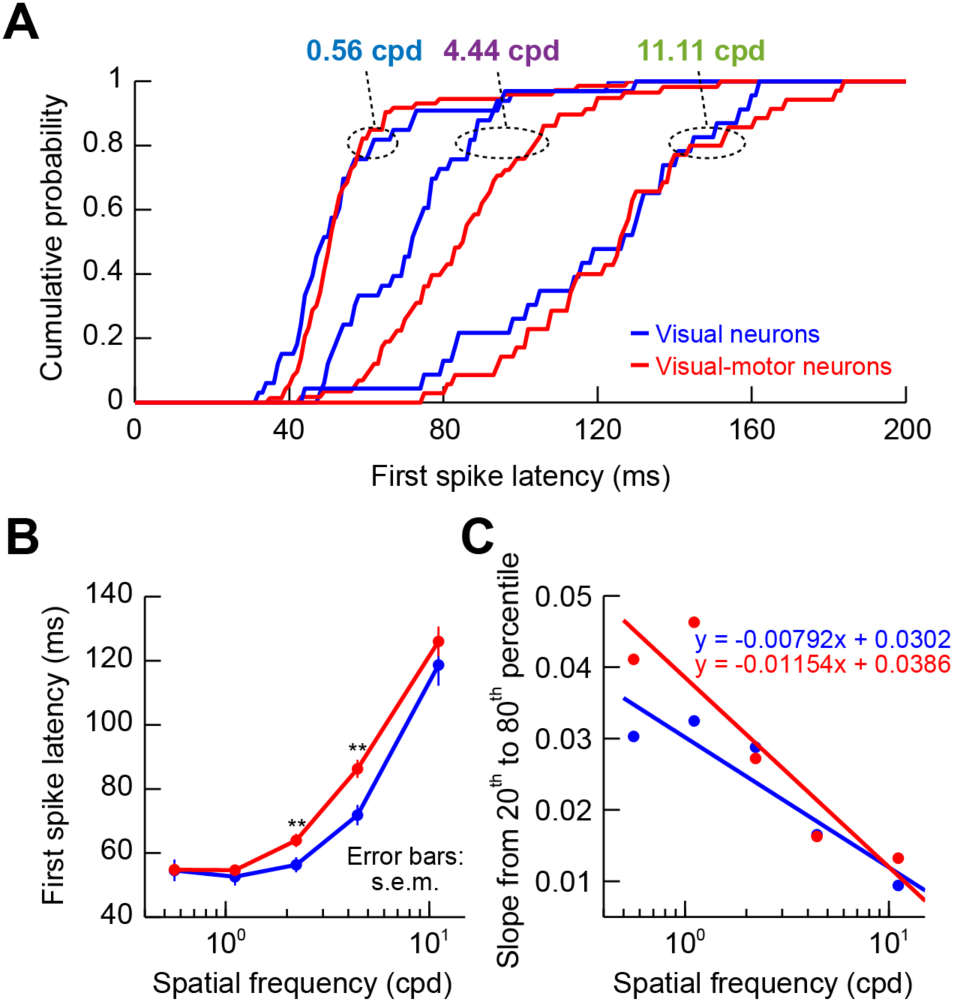
Response latencies of visual and visual-motor SC neurons. (**A**) Cumulative histograms of first-spike latency as in Fig. 4, but after separating neurons as either being purely visual (blue) or visual-motor (red). To reduce clutter, only 3 spatial frequencies are shown. As can be seen, the dependence of response latency on spatial frequency was similar whether neurons were purely visual or visual-motor, but visual neurons tended to exhibit slightly shorter latencies, especially at 4.44 cpd. (**B**) Mean response latencies across neurons as a function of spatial frequency, again as in Fig. 4, but separating visual and visual-motor neurons. Consistent with **A**, response latency increased with increasing spatial frequency for both types of neurons. Also, again consistent with **A**, visual neurons showed earlier responses than visual-motor neurons, especially for intermediate spatial frequencies. Asterisks imply p<0.01 for the comparison between visual and visual-motor neurons at a given spatial frequency (Ranksum test). (**C**) Estimate of variance in response latency as a function of spatial frequency, as in Fig. 4. Both visual and visual-motor neurons behaved similarly as a function of spatial frequency, with higher variability of response latencies across neurons for higher spatial frequencies.

**Figure 4-Figure Supplement 2.**
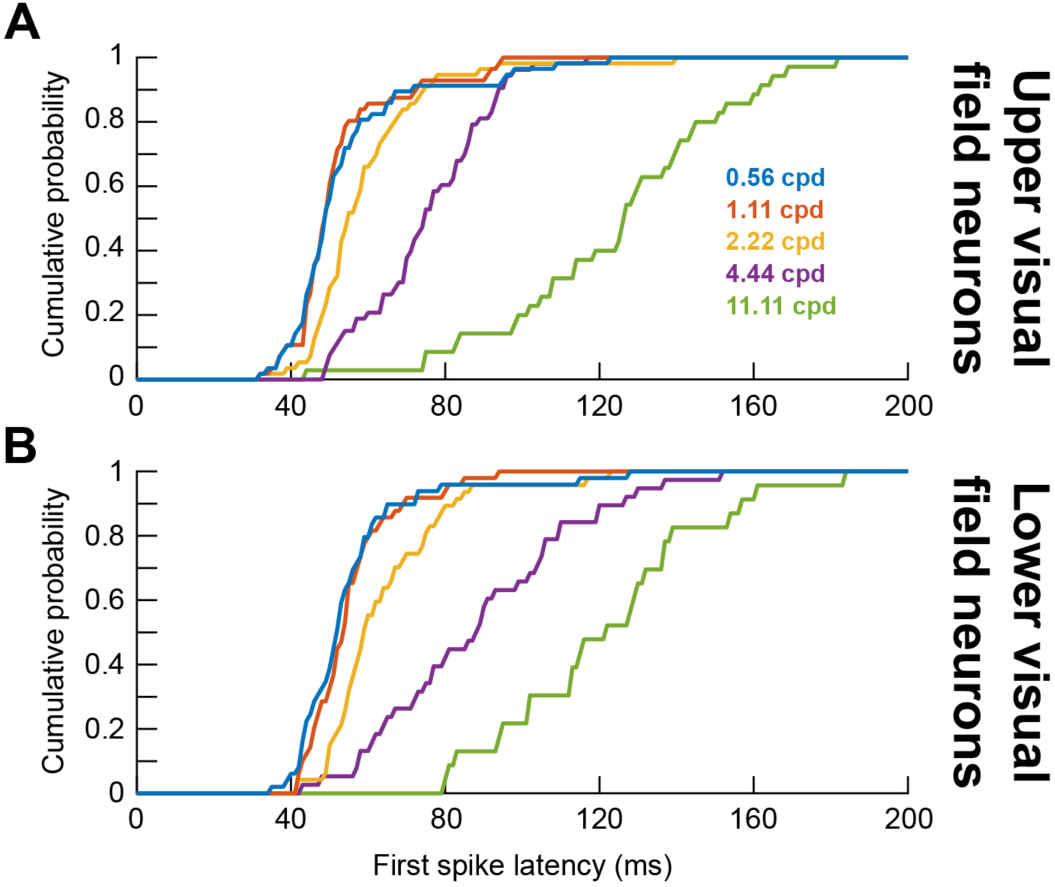
Response latencies for upper and lower visual field neurons. (**A**) Same as Fig. 4A but for neurons in the upper visual field representation of the SC. (**B**) Same as Fig. 4A but for neurons in the lower visual field representation of the SC. In both cases, the rank ordering of response latencies as a function of spatial frequency is evident, with an additional observation of upper visual field neurons responding faster in general than lower visual field neurons (Hafed and Chen, 2016).

